# Molecular Modeling Evaluation of the Binding Effect of Ritonavir, Lopinavir and Darunavir to Severe Acute Respiratory Syndrome Coronavirus 2 Proteases

**DOI:** 10.1101/2020.01.31.929695

**Authors:** Shen Lin, Runnan Shen, Jingdong He, Xinhao Li, Xushun Guo

## Abstract

Three anti-HIV drugs, ritonavir, lopinavir and darunavir, might have therapeutic effect on coronavirus disease 2019 (COVID-19). In this study, the structure models of two severe acute respiratory syndrome coronavirus 2 (SARS-CoV-2) proteases, coronavirus endopeptidase C30 (CEP_C30) and papain like viral protease (PLVP), were built by homology modeling. Ritonavir, lopinavir and darunavir were then docked to the models, respectively, followed by energy minimization of the protease-drug complexes. In the simulations, ritonavir can bind to CEP_C30 most suitably, and induce significant conformation changes of CEP_C30; lopinavir can also bind to CEP_C30 suitably, and induce significant conformation changes of CEP_C30; darunavir can bind to PLVP suitably with slight conformation changes of PLVP. It is suggested that the therapeutic effect of ritonavir and lopinavir on COVID-19 may be mainly due to their inhibitory effect on CEP_C30, while ritonavir may have stronger efficacy; the inhibitory effect of darunavir on SARS-CoV-2 and its potential therapeutic effect may be mainly due to its inhibitory effect on PLVP.

## 1. Introduction

In December 2019, coronavirus disease 2019 (COVID-19) caused by severe acute respiratory syndrome coronavirus 2 (SARS-CoV-2) occurred in Wuhan, China, and then quickly became a rising global concern since World Health Organization had declared it as the sixth public health emergency of international concern (PHEIC) on 30 January 2020. By the time of 14:50 on February 8, 2020, 34638 COVID-19 cases had been confirmed in China and 723 patients died from it, resulting in a mortality of 2.2%. SARS-CoV-2 is a novel species of non-segmented positive-sense RNA virus which can spread and distribute in humans and other mammals [1]. Previous studies have discovered that SARS-CoV-2 belongs to a distinct clade from the β-coronavirus which is associated with human severe acute respiratory syndrome (SARS) and Middle East respiratory syndrome (MERS) [2]. Shi ZL et al found that SARS-CoV-2 shares 79.5% sequence identity to SARS-related coronavirus and is 96% identical at the whole-genome level to a bat coronavirus [3], which corresponds to that the origin may be wild animals sold in Huanan Seafood Wholesale Market [4]. Recent clinical research found that dyspnea might occur on the fifth day and acute respiratory distress syndrome on the eighth day from first symptom of COVID-19 in median, organ dysfunction and death can also occur in severe COVID-19 cases [5, 6]. However, there is still no specific treatment for COVID-19. Finding drugs that can fight against SARS-CoV-2 is a top priority.

The trial fifth edition of diagnosis and treatment guideline of COVID-19 issued by National Health Commission of the People’s Republic of China (http://www.nhc.gov.cn/) recommends to use Kaletra for treatment. Kaletra, an anti-HIV drug which is composed of two protease inhibitors, ritonavir (CAS#: 155213-67-5) and lopinavir (CAS#: 192725-17-0), might have therapeutic effect on coronavirus diseases like SARS and MERS [7–10]. However, whether it can inhibit SARS-CoV-2 or treat COVID-19 lacks clinical evidences and randomized clinical trials, and the safety of its use in COVID-19 patients is unclear. Otherwise, Lanjuan Li, infectious disease scientist, academician of Chinese Academy of Engineering, recommended darunavir (CAS#: 206361-99-1), also an HIV protease inhibitor, as a treatment for COVID-19. Although the inhibitory effect of darunavir on SARS-CoV-2 has been verified in vitro, its therapeutic effect on COVID-19 is still unknown. At the same time, the mechanism of how these drugs inhibit SARS-CoV-2 is also unknown.

As coronaviruses, including SARS-CoV-2, synthesize polyproteins followed by hydrolyzed to produce their structure and function proteins [11–13], it is suggested that ritonavir, lopinavir and darunavir may block the multiplication cycle of SARS-CoV-2 by inhibiting its proteases. To preliminarily understand the inhibitory effects of the drugs, in this study, molecular models of the proteases were built by homology modeling, followed by docking the drugs to the proteases, respectively. In addition, the dynamic interactions between the drugs and the protease were also simulated to evaluate the binding effects of the drugs on the proteases.

## 2. Materials and Methods

### 2.1 Homology Modeling of the Proteases

Since there was no gene of protease identified in the ten SARS-CoV-2 genes directly, the protease-like conserved domains in the orf1ab polyprotein (GenBank: QHO60603.1) of the virus were analyzed through NCBI Conserved Domain Search Service (https://www.ncbi.nlm.nih.gov/Structure/cdd/wrpsb.cgi) [14–17]. The sequences of the found protease-like domains were then used to build molecular models by homology modeling through SWISS-MODEL (https://swissmodel.expasy.org/) [18–22]. To evaluate qualities, Verify score, Errat score, Prove score and Ramachandran plots of the built models were automatically created by Structure Analysis and Verification server (version 5.0, https://servicesn.mbi.ucla.edu/SAVES/).

### 2.2 Molecular Docking of the Drugs

To dock ritonavir, lopinavir and darunavir to the proteases, respectively, Discovery Studio software (version 2.5, Accelrys Software Inc.) was used. While docking, one of the proteases would be set as receptor, and all its possible docking sites would be found. A sphere whose radius was 17 units would then be set at the optimal docking site, followed by semi-flexible docking of unprotonated ritonavir, lopinavir and darunavir to the receptor protease through Dock Ligands (Libdock) module of Discovery Studio software, respectively. The preliminary docking results would be evaluated by Libdock scores and intermolecular interactions through Discovery Studio software.

To analyze the structure changes of proteases caused by induced-fit effect after binding the drugs, Steepest Descent and Conjugate Gradient Algorithms were used to minimize the energy of the protease-drug complexes successively through Minimization module of Discovery Studio software. During calculation, fixed constraint of the backbone of proteins was used first and removed later.

## 3. Results

### 3.1 Molecular Models of the Proteases

By searching the conserved domains of the sequence of orf1ab polyprotein, two protease-like domains were found. One domain is coronavirus endopeptidase C30 (CEP_C30), located from 3292 to 3569 of the polyprotein sequence; the other domain is papain like viral protease (PLVP), located from 1564 to 1880 of the sequence. The sequence of PLVP was directly used for subsequent homology modeling, while the 5 residues at the C-terminal of the sequence of CEP_C30 were deleted before subsequent homology modeling to minimize disordered structures.

While modeling CEP_C30, 8 models were built according to 8 automatically found templates. The top-rated model in the 8 models is a homo-dimer with no ligand, and was selected to represent the structure of CEP_C30. While modeling PLVP, 29 models were built according to 16 automatically found templates. The top-rated model in the 29 models is a monomer with a zinc ion ligand, and was selected to represent the structure of PLVP. The structures of the two selected models are available in Supplemental Files S1 and S2. The detailed information, evaluation scores and Ramachandran plots of the two models are shown in Figure 1.

**Figure 1.**
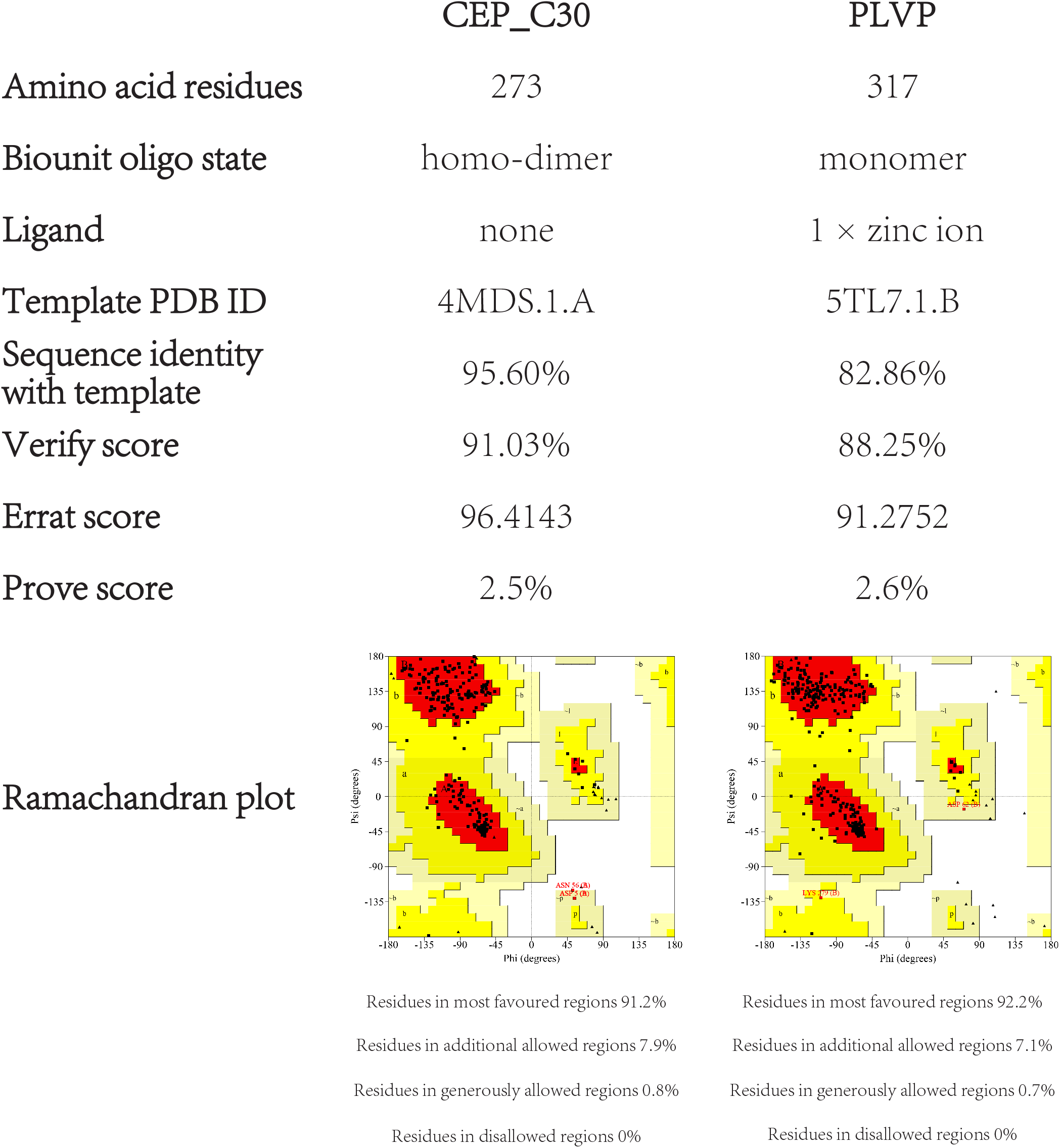
The detailed information, evaluation scores and Ramachandran plots of structure models of CEP_C30 and PLVP. The structure model of CEP_C30 is a homo-dimer with no ligand, built by taking PDB: 4MDS.1.A as template. Each chain of the model of CEP_C30 is consist of 273 amino acid residues, with 95.60% sequence identity with the template. The structure model of PLVP is a monomer with a zinc ion ligand, built by taking PDB: 5TL7.1.B as template. The model of PLVP is consist of 317 amino acid residues, with 82.86% sequence identity with the template. The evaluation scores and Ramachandran plots of two models were automatically created by Structure Analysis and Verification server (version 5.0, https://servicesn.mbi.ucla.edu/SAVES/). There are three scores used to evaluate the models: Verify score, Errat score and Prove score. The higher Verify score and Errat score are, as well as the lower Prove score is, the more reasonably the model is built.

### 3.2 Molecular Docking of the Drugs

The optimal docking site of CEP_C30 is a pocket between its two subunits. Through docking drugs to this site, 100 poses were found when docking ritonavir, with the Libdock score of the optimal pose 192.346; 88 poses were found when docking lopinavir, with the Libdock score of the optimal pose 147.123; and 49 poses were found when docking darunavir, with the Libdock score of the optimal pose 149.404. The optimal docking site of PLVP is a pocket in one side of the protein. Through docking drugs to this site, 4 poses were found when docking ritonavir, with the Libdock score of the optimal pose 164.153; 3 poses were found when docking lopinavir, with the Libdock score of the optimal pose 107.137; and 8 poses were found when docking darunavir, with the Libdock score of the optimal pose 139.543. The interactions between the proteases and drugs at the optimal poses are shown in Figure 2, and the structures of the docked protease-drug complexes at the optimal pose before energy minimization are available in Supplemental Files S3 to S8.

**Figure 2.**
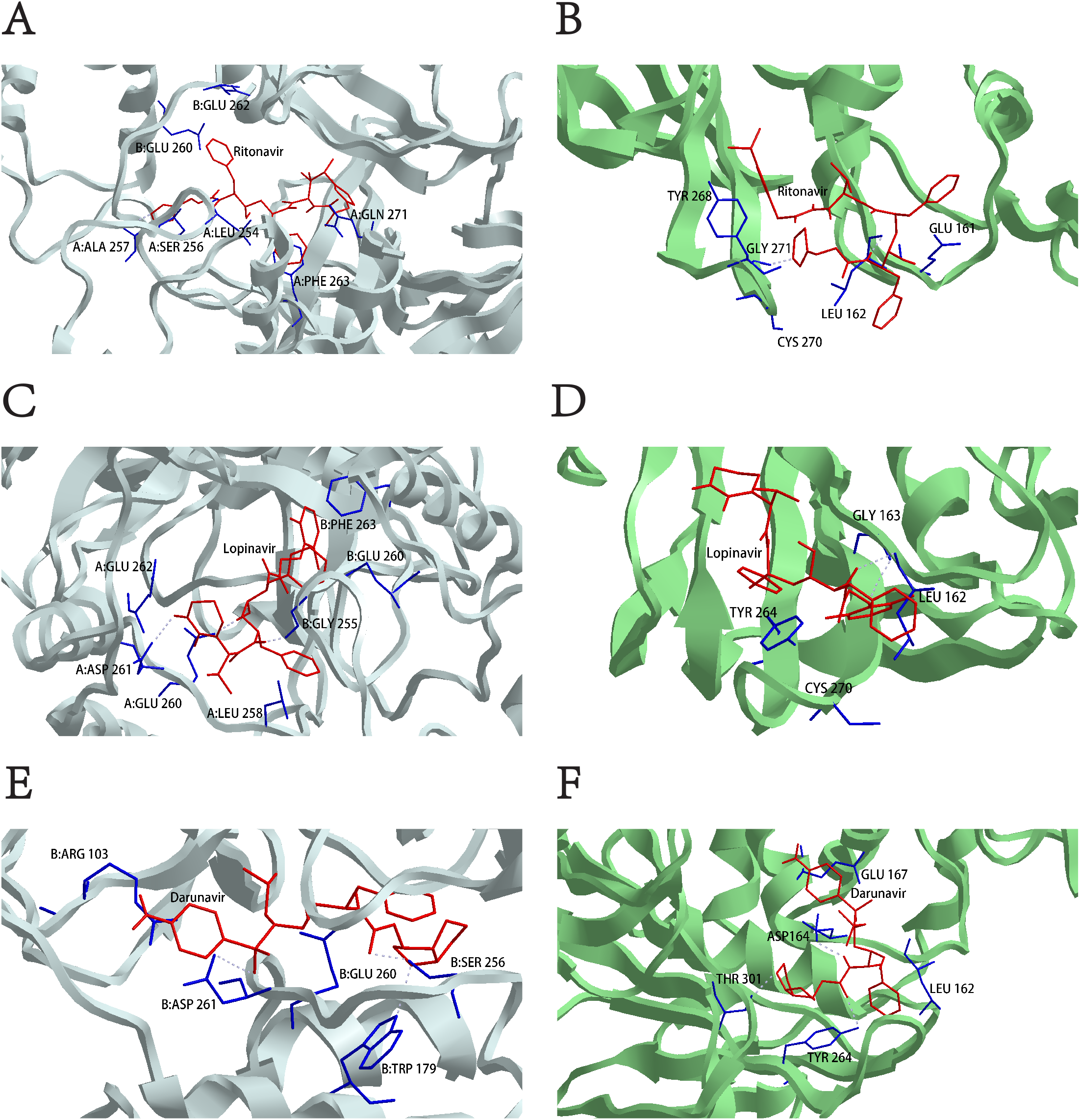
The interactions between the proteases and drugs at the optimal poses before energy minimization. The interactions (A) between ritonavir and CEP_C30; (B) between ritonavir and PLVP; (C) between lopinavir and CEP_C30; (D) between lopinavir and PLVP; (E) between darunavir and CEP_C30; and (F) between darunavir and PLVP before energy minimization. The main chains of CEP_C30 are shown in white, and of PLVP are shown in green. The side chains of CEP_C30 and PLVP which interact with ritonavir, lopinavir or darunavir are shown in blue. Ritonavir, lopinavir and darunavir are shown in red. The hydrogen bonds are shown in white interrupted lines.

Through energy minimization, all protease-drug complexes had some structure changes. The differences between before and after energy minimization of protease-drug complexes are shown in Figure 3, and the structures of protease-drug complexes after energy minimization are available in Supplemental Files S9 to S14.

**Figure 3.**
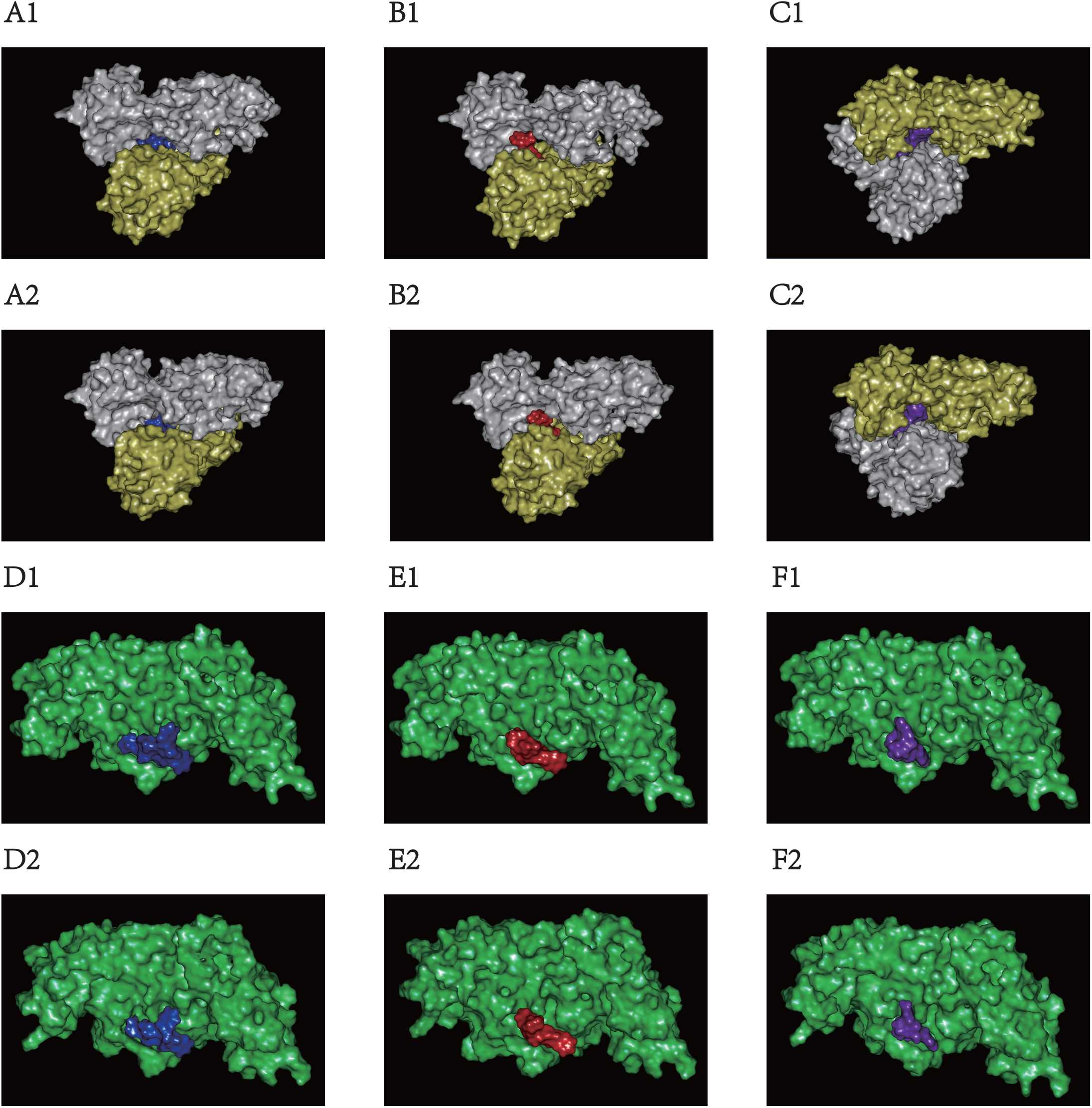
The differences between before and after energy minimization of protease-drug complexes. The structures of (A1) CEP_C30-ritonavir complex before energy minimization; (A2) CEP_C30-ritonavir complex after energy minimization; (B1) CEP_C30-lopinavir complex before energy minimization; (B2) CEP_C30-lopinavir complex after energy minimization; (C1) CEP_C30-darunavir complex before energy minimization; (C2) CEP_C30-darunavir complex after energy minimization; (D1) PLVP-ritonavir complex before energy minimization; (D2) PLVP-ritonavir complex after energy minimization; (E1) PLVP-lopinavir complex before energy minimization; (E2) PLVP-lopinavir complex after energy minimization; (F1) PLVP-darunavir complex before energy minimization; and (F2) PLVP-darunavir complex after energy minimization. The chain A of CEP_C30 is shown in white, and chain B of CEP_C30 is shown in yellow. The main chain of PLVP is shown in green. Ritonavir is shown in blue, lopinavir in red, and darunavir in purple.

## 4. Discussion

### 4.1 The Significance of the Molecular Models

In this study, two models of two protease-like domains, CEP_C30 and PLVP, of orf1ab polyprotein of SARS-CoV-2 were built. Since these two models were built by taking highly homologous crystal structures as templates, their qualities are quite considerable. As it is difficult to obtain the crystal structure of CEP_C30 and PLVP in a short time, there is great significance to use these reliable structure models to predict their dynamic interactions with other molecules.

### 4.2 The Binding Effects of the Drugs

For ritonavir and lopinavir, these two drugs seem more suitable to bind to CEP_C30 rather than PLVP. Through energy minimization, it is suggested that after docking ritonavir or lopinavir, CEP_C30 can be induced to change its conformation significantly to bind the drug tightly. Moreover, ritonavir is more suitable than lopinavir to bind to CEP_C30. As shown in Figure 2 and 3, there are more interactions between CEP_C30 and ritonavir, and the binding between them is tighter. On the contrary, neither ritonavir nor lopinavir seems suitable to bind to PLVP, since the bindings become less tight after energy minimization.

For darunavir, this drug seems more suitable to bind to PLVP rather than CEP_C30. Through energy minimization, it is suggested that after docking to PLVP, darunavir can change its conformation significantly to fit PLVP, whose conformation will also be induced to slightly change. On the contrary, darunavir seems not suitable to bind to CEP_C30, since the binding between darunavir and CEP_C30 becomes less tight after energy minimization.

### 4.3 The Possible Inhibitory Mechanisms of the Drugs

Since the catalytic mechanisms of CEP_C30 and PLVP domains are both unknown, it is unable to determine whether ritonavir, lopinavir and darunavir are competitive or non-competitive inhibitors, although these drugs are all peptide analogues and seem to be competitive.

As the binding of ritonavir and lopinavir might lead to significant conformation changes of CEP_C30, there is no conclusion whether ritonavir and lopinavir are competitive or non-competitive, since both competitive and non-competitive inhibitors may significantly change the conformation of receptor. For darunavir, as it can bind to PLVP suitably by changing its conformation with slight conformation changes of PLVP, it is more likely to be a competitive inhibitor of PLVP.

In conclusion, the therapeutic effect of ritonavir and lopinavir on COVID-19 and other coronavirus disease may be mainly due to their inhibitory effect on CEP_C30, while ritonavir may have stronger efficacy; the inhibitory effect of darunavir on SARS-CoV-2 and its potential therapeutic effect may be mainly due to its inhibitory effect on PLVP. However, there are still limitations in this study. As the catalytic mechanisms of CEP_C30 and PLVP are still unknown, in further studies, it should still be focused on to figure out these mechanisms and how ritonavir, lopinavir and darunavir block these procedures. The safety and actual effects of these drugs on human body should also be carried out and verified by randomized clinical control studies further.

## Supplemental Materials

Supplemental File S1. The structure model of CEP_C30.

Supplemental File S2. The structure model of PLVP.

Supplemental File S3. The structures of docked CEP_C30-ritonavir complex at the optimal pose before energy minimization.

Supplemental File S4. The structures of docked CEP_C30-lopinavir complex at the optimal pose before energy minimization.

Supplemental File S5. The structures of docked CEP_C30-darunavir complex at the optimal pose before energy minimization.

Supplemental File S6. The structures of docked PLVP-ritonavir complex at the optimal pose before energy minimization.

Supplemental File S7. The structures of docked PLVP-lopinavir complex at the optimal pose before energy minimization.

Supplemental File S8. The structures of docked PLVP-darunavir complex at the optimal pose before energy minimization.

Supplemental File S9. The structures of docked CEP_C30-ritonavir complex after energy minimization.

Supplemental File S10. The structures of docked CEP_C30-lopinavir complex after energy minimization.

Supplemental File S11. The structures of docked CEP_C30-darunavir complex after energy minimization.

Supplemental File S12. The structures of docked PLVP-ritonavir complex after energy minimization.

Supplemental File S13. The structures of docked PLVP-lopinavir complex after energy minimization.

Supplemental File S14. The structures of docked PLVP-darunavir complex after energy minimization.

## Acknowledgments

The authors thank Haibin Luo (School of Pharmaceutical Sciences, Sun Yat-sen University, Guangzhou, China) for providing the Discovery Studio software, and thank Wufei Wang (School of Aeronautics and Astronautics, University of Electronic Science and Technology of China, Chengdu, China) for proofreading the manuscript.

## Declarations

There is no conflict of interest to declare.

